## Supplemental Information (Figures S1-S10,Table S1) for "Semi-field and surveillance data define the natural diapause timeline for *Culex pipiens* across the United States"

### Supplemental Figures

**Figure S1.** Detection of autogenous mosquitoes in the *Culex pipiens* lab colony.

**Figure S2.** Description of sites used for semi-field studies of mosquito diapause.

**Figure S3.** Temperature and photoperiod associated with semi-field experimental cohorts.

**Figure S4.** Evaluating pupal exposure to environmental signals under semi-field conditions.

**Figure S5.** Temperature comparisons of 2020 and 2021 to 30-year averages.

**Figure S6.** Seasonal changes to diurnal temperature fluctuations.

**Figure S7.** Mosquito surveillance sites in Iowa.

**Figure S8.** Gravid populations are distinct from general *Cx. pipiens* mosquito abundance.

**Figure S9.** Geographic locations and elevations of trapping site locations across the United States.

**Figure S10.** End-of-season temperature and photoperiod conditions at trapping sites across the United States.

### Supplemental Tables

**Table S1:** Larval Culex pipiens semi-field group descriptions, Ames IA.

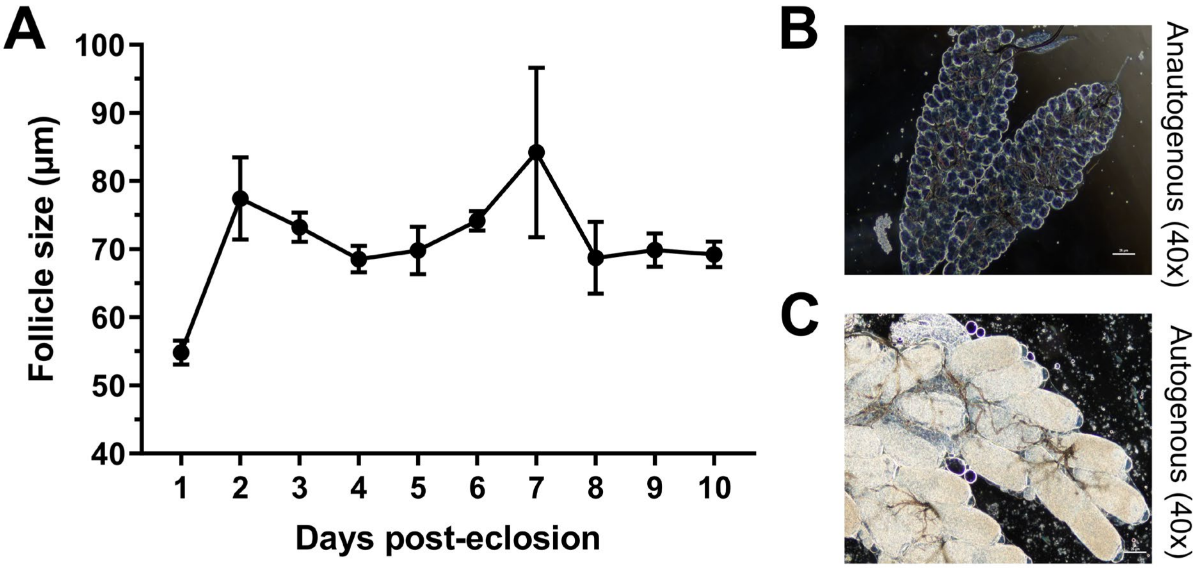

**Figure S1. Detection of autogenous mosquitoes in the *Culex pipiens* lab colony.** Adult female *Cx. pipiens* were dissected daily from 1 to 10 days post-eclosion to examine ovarian development and average follicle size (**A**). A total of 10 mosquitoes were used to calculate the average follicle size for days 1-4, while 4 mosquitoes were used for days 5-10. Example images display “normal” anautogenous ovaries (**B**) or less frequent examples of autogenous mosquito ovaries (**C**) as evidenced by the elongation and granulation of the follicle in the absence of a blood meal. Images are displayed with 40x magnification.

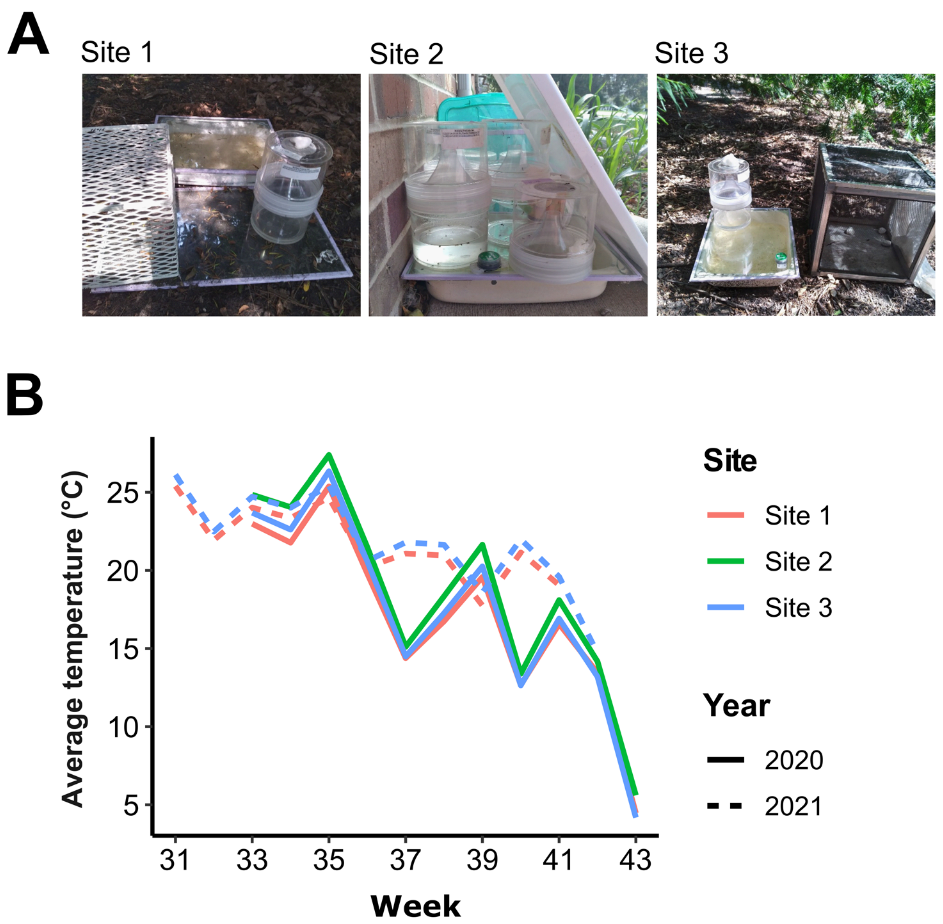

**Figure S2. Descriptions and temperatures of sites used in semi-field studies.** Three site locations were selected in 2020 in Ames, IA (USA) for semi-field studies to examine mosquito diapause induction (**A**). Sites 1 and 3 were used again in 2021 for the second year of the study. (**B**) Temperatures were recorded at each site using Hobo MX2202 data loggers to evaluate any differences in temperature in the microenvironment.

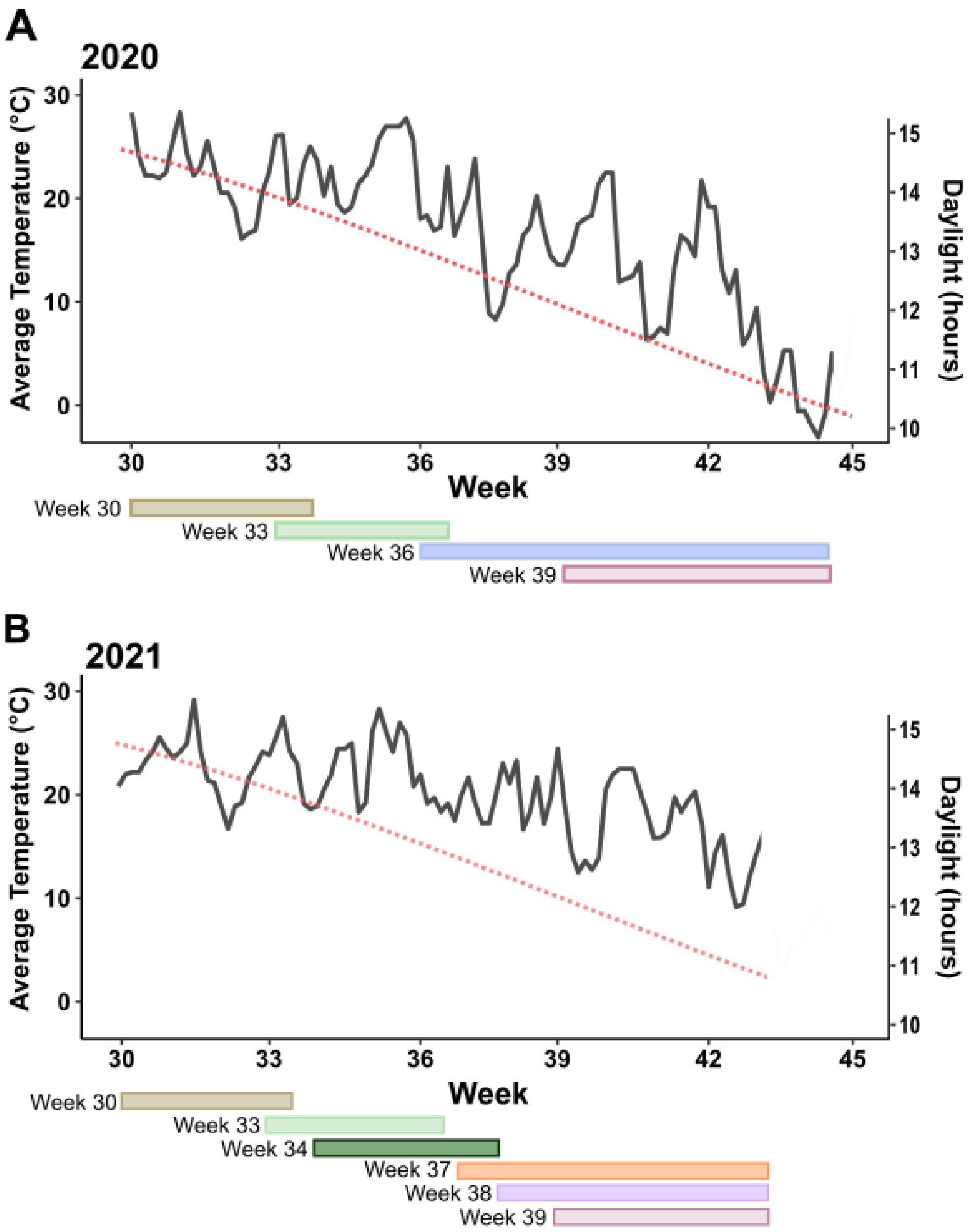

**Figure S3. Temperature and photoperiod associated with semi-field experimental cohorts.** Temperature (solid line) and photoperiod (dotted line) values are displayed for the experimental cohorts used in semi-field experiments for 2020 (**A**) and 2021 (**B**). For each year of experiments, larval cohorts are denoted below with the time of field placement (week) to the last adult eclosion or death. The length of bar represents the time in which each cohort was active.

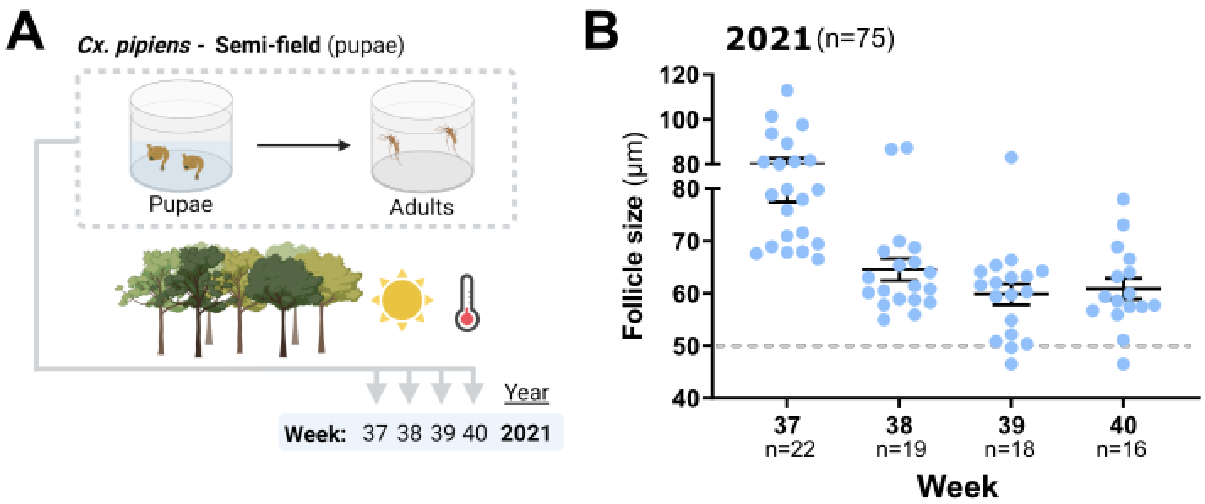

**Figure S4. Evaluating pupal exposure to environmental signals under semi-field conditions.** To examine if laboratory-reared pupae could undergo emergence in diapause under semi-field conditions, pupae were placed in "mosquito breeder" containers in semi-field conditions during timepoints where larval cohorts were receptive to diapause (**A**). A total of 75 mosquitoes were examined to determine if they emerged in reproductive diapause (**B**). Each dot represents the average follicle size for an individual mosquito, with a 50 µm threshold (dotted line) used to determine individuals in the diapause state. The mean follicle size (+/- SEM) is displayed for each experimental cohort. n, number of individual mosquitoes examined.

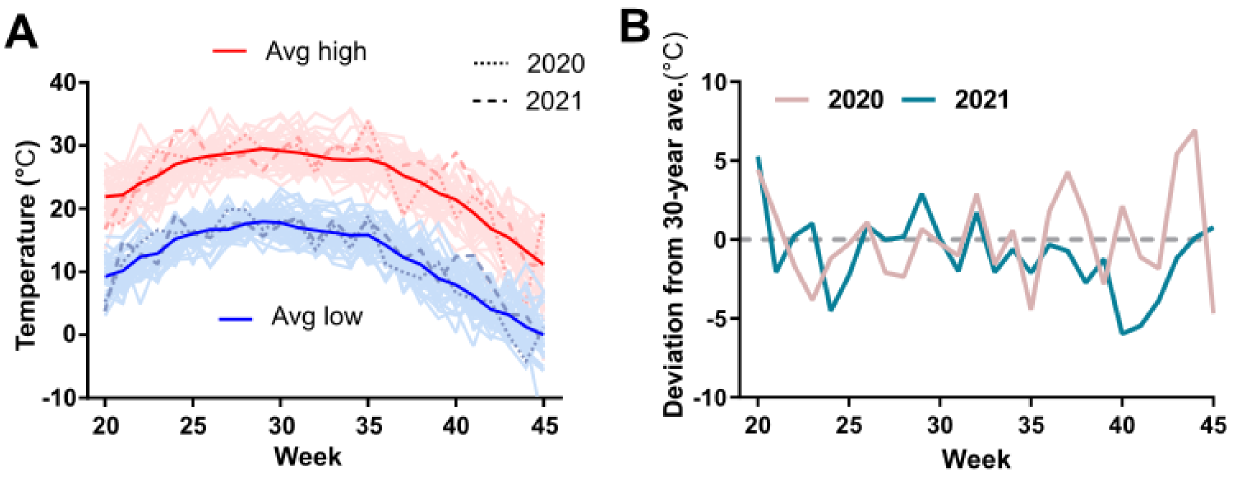

**Figure S5. Temperature comparisons of 2020 and 2021 to 30-year averages.** To place the temperatures of our two-year (2020 and 2021) semi-field study in context, average high and low temperatures were compared to the historical thirty-year average weekly high and low temperatures in Central Iowa (**A**). The thirty-year average high (red) and low (blue) temperatures are displayed by week, with the average temperatures for individual years displayed by light red (high) or light blue (low) lines. or presented in light red (high) and light blue (low). Average trends for each variable over all years are presented in dark red and dark blue, respectively. The values for 2020 and 2021 are respectfully displayed by dotted or marked lines. (**B**) The deviation in average temperature for 2020 and 2021 is displayed by week from the thirty-year average.

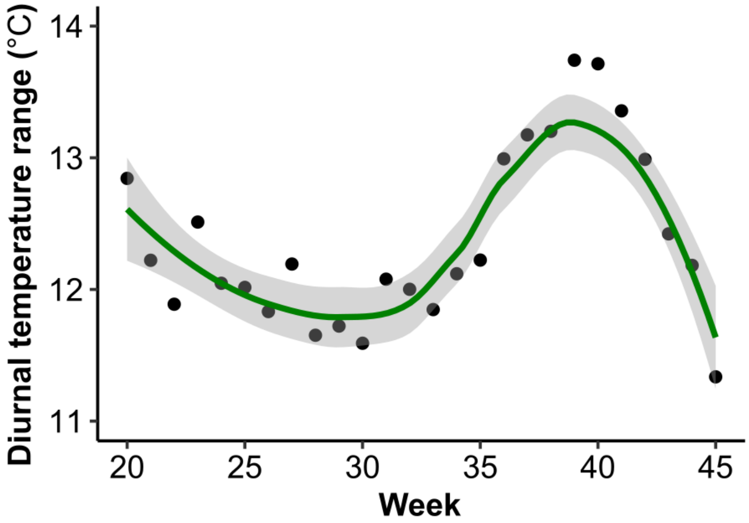

**Figure S6. Seasonal changes to diurnal temperature fluctuations.** The diurnal temperature range (difference between high and low temperatures) is displayed by week.

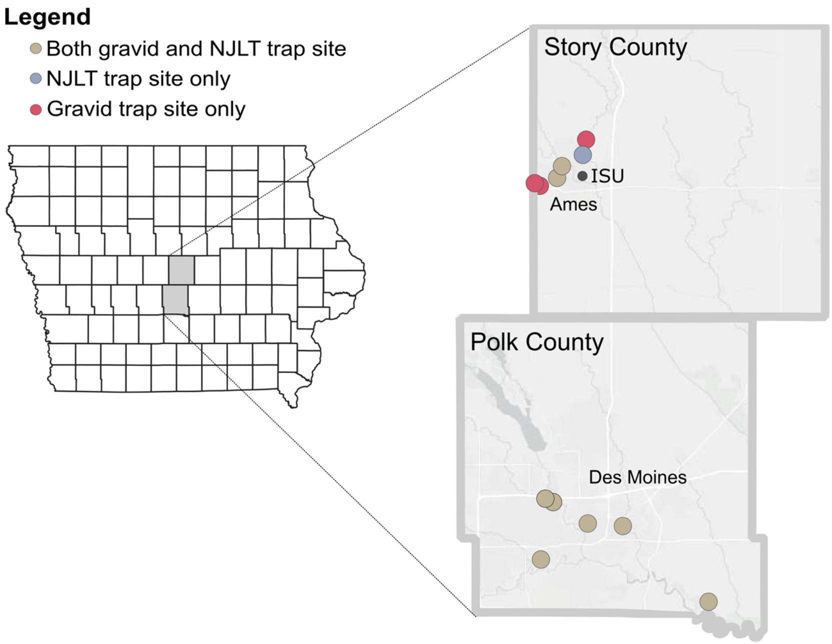

**Figure S7. Mosquito surveillance sites in Iowa.** Two types of mosquito trapping data were used for this study: Gravid (captures reproductive trends) and New Jersey Light Trap (NJLT; captures general trends). A total of 16 sites were included for their gravid records (Gravid-only sites in red), and 14 sites for their NJLT records (NJLT-only sites in blue) from 2016-2021. The sites denoted in tan provided both gravid and NJLT data.

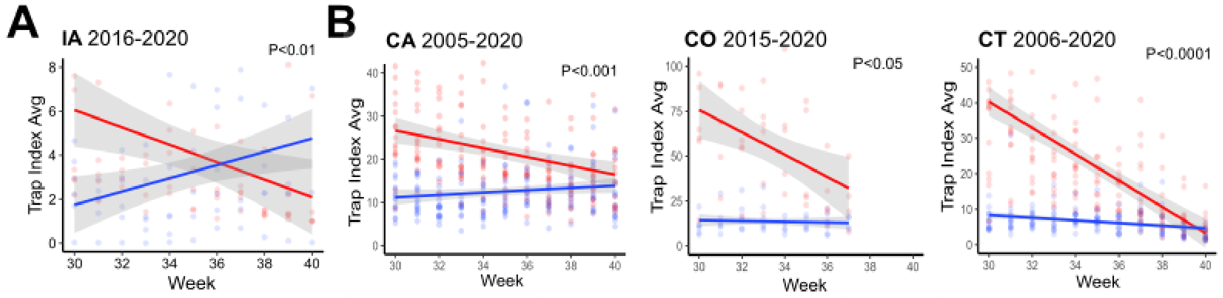

**Figure 8. Gravid populations are distinct from general *Cx. pipiens* mosquito abundance.** Comparisons of end-of-season gravid trap data (reproductive populations) and New Jersey Light Trap (NJLT; general abundance) for *Cx. pipiens* group mosquitoes in Iowa (**A**) and other locations across the country (**B**). For both **A** and **B**, data display the normalized trap index values from weeks 30 to 40 for the years shown above each graph. Average weekly gravid or NJLT values across all years of trapping are shown by the solid lines (95% confidence intervals in grey), with data from individual years displayed for gravid (light red) or NJLT (light blue) tapping data. The annual slope values for gravid and NJLT trends were compared using a linear regression, with significance indicated for each location.

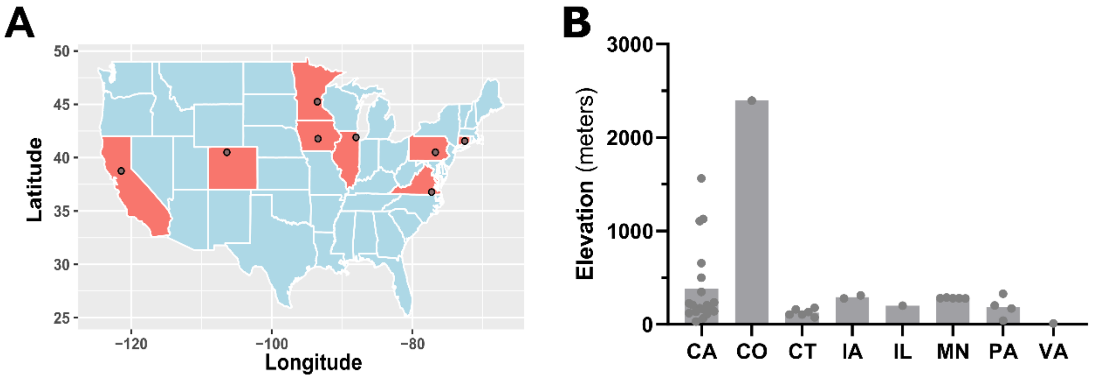

**Figure S9. Geographic locations and elevations of trapping site locations across the United States.** (**A**) The geographic location (latitude and longitude) of each trapping site location is displayed on the map, with the approximate site location (or centroid of the trapping area) denoted by a circle. (**B**) The average elevation of the participating counties in each state included in our analysis, with individual counties displayed by the grey dots.

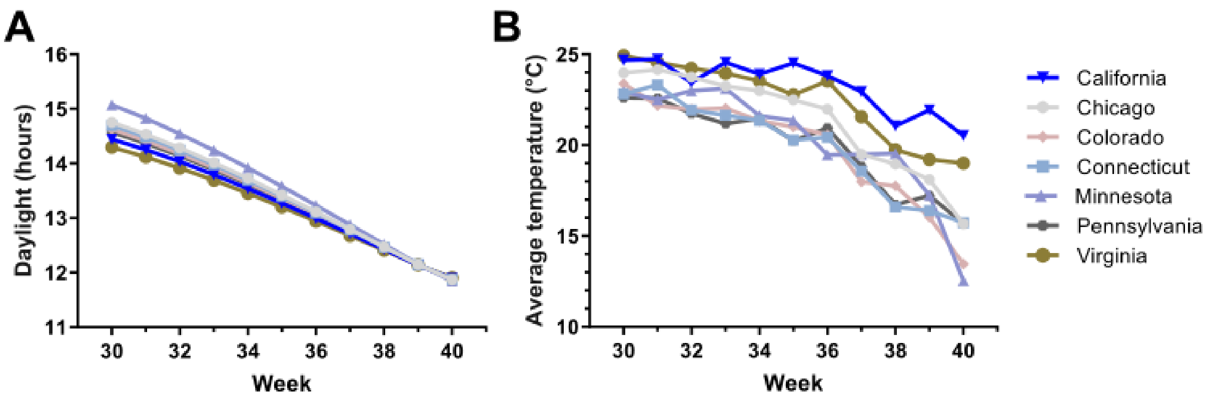

**Figure S10. End-of-season temperature and photoperiod conditions at trapping sites across the United States.** The average photoperiod (measured as hours of daylight) (**A**) or average temperature (**B**) for each of the locations across the United States.

**Table S1**: Larval *Culex pipiens* semi-field group descriptions, Ames IA.

| Year | Start Date | Week Start | Week of first pupae | Week End | Start Temp | Temp at first pupae | End Temp | Start Photo | Photo at first pupae | End Photo | Diapause recorded? |
|---|---|---|---|---|---|---|---|---|---|---|---|
| 2020 | July-19 | 30 | 31 | 33 | 23.7 | 19.5 | 23.5 | 14.7 | 14.5 | 14.0 | N |
| 2020 | Aug-9 | 33 | 34 | 36 | 23.5 | 21.1 | 18.7 | 14.0 | 13.7 | 13.1 | N |
| 2020 | Aug-20 | 36 | 38 | 43 | 18.7 | 16.2 | 3.8 | 13.1 | 12.4 | 10.8 | Y |
| 2020 | Sep-20 | 39 | 41 | 43 | 18.5 | 13.8 | 3.8 | 12.1 | 11.4 | 10.8 | Y |
| 2021 | Jul-25 | 30 | 31 | 34 | 23.2 | 24.9 | 22.3 | 14.7 | 14.5 | 13.7 | N |
| 2021 | Aug-15 | 33 | 34 | 35 | 24.0 | 22.3 | 23.9 | 14.0 | 13.7 | 13.4 | N |
| 2021 | Aug-22 | 34 | 35 | 36 | 22.3 | 23.9 | 20.9 | 13.7 | 13.4 | 12.8 | Y |
| 2021 | Sep-12 | 37 | 39 | 42 | 19.4 | 16.9 | 14.2 | 12.8 | 12.1 | 11.2 | Y |
| 2021 | Sep-18 | 38 | 39 | 42 | 20.3 | 16.9 | 14.2 | 12.5 | 12.1 | 11.2 | Y |
| 2021 | Sep-25 | 39 | 41 | 42 | 16.9 | 11.5 | 14.2 | 12.1 | 11.5 | 11.2 | Y |
